# Hyperactivation of YAP/TAZ drives alterations in mesangial cells through stabilization of N-MYC in diabetic nephropathy

**DOI:** 10.1101/2022.01.29.478286

**Authors:** Seunghyeok Choi, Sang Heon Suh, Hosung Bae, Kyung Pyo Kang, Hyuek Jong Lee, Gou Young Koh

**Affiliations:** Graduate School of Medical Science and Engineering, Korea Advanced Institute of Science and Technology, Daejeon 34141, Republic of Korea; Center for Vascular Research, Institute for Basic Science, Daejeon 34141, Republic of Korea; Department of Internal Medicine, Research Institute of Clinical Medicine, Jeonbuk National University Medical School, Jeonju 54907, Republic of Korea; Biomedical Research Institute, Jeonbuk National University Hospital, Jeonju 54907, Republic of Korea

## Abstract

Mesangial cells (MCs) in the kidney are central to maintaining glomerular integrity, and their impairment leads to major glomerular diseases including diabetic nephropathy (DN). Although high blood glucose elicits abnormal alterations in MCs, the underlying molecular mechanism is poorly understood. Here, we show that YAP and TAZ, the final effectors of the Hippo pathway, are highly increased in MCs of patients with DN and of Zucker diabetic fatty rats. Moreover, high glucose directly induces activation of YAP/TAZ through the canonical Hippo pathway in cultured MCs. Hyperactivation of YAP/TAZ in mouse model MCs recapitulates the hallmarks of DN, including excessive proliferation of MCs and extracellular matrix deposition, endothelial cell impairment, glomerular sclerosis, albuminuria, and reduced glomerular filtration rate. Mechanistically, activated YAP/TAZ bind and stabilize N-Myc protein, one of the Myc family of oncogenes. N-Myc stabilization leads to aberrant enhancement of its transcriptional activity and eventually to MC impairments and DN pathogenesis. Together, these findings shed light on how high blood glucose in diabetes mellitus leads to DN and support a rationale that lowering blood glucose in diabetes mellitus could delay DN pathogenesis.

## Introduction

Mesangial cells (MCs) are specialized microvascular pericytes with a central role in maintaining the structure and function of the renal glomerulus (1–4). Because MCs constitute the central portion of the glomerulus, they both provide structural support of the glomerular capillary loop and regulate glomerular filtration using their contractile properties (1, 2, 4, 5). Dynamic and versatile interactions among MCs, glomerular endothelial cells (ECs) and podocytes are key to maintaining glomerular integrity and in recovery after injury (1, 2, 4, 6–9). For these reasons, aberrant alterations in MCs are involved in a wide variety of glomerular diseases (1, 2, 4, 7, 8), particularly in immune-mediated and metabolic renal diseases (8, 10). These alterations in MCs are critically involved in reduced renal function in diabetic nephropathy (DN) (11–13). Continuous exposure to high blood glucose elicits structural and functional alterations in glomerular ECs and MCs, including disrupted EC fenestrations, MC proliferation, excessive accumulation of extracellular matrix (ECM) around MCs, and reduced glomerular filtration (12, 13). Nevertheless, an exact and detailed underlying molecular mechanism for how high blood glucose induces these alterations has yet to be elucidated.

The mammalian Hippo pathway regulates cellular differentiation and proliferation, tissue homeostasis, and stem cell renewal (14–16). The key components of the cascade are large tumor suppressor 1/2 (LATS1/2) and the final transcriptional regulators, Yes-associated protein (YAP) and transcriptional coactivator with a PDZ-binding motif (TAZ), which primarily bind to TEA domain transcription factor (TEAD) to exert their functions (14–16). The Hippo pathway is crucial in embryonic kidney development (17, 18). Silencing YAP in early embryonic nephron progenitor cells leads to hypoplastic kidney, and TAZ knockout mice show impairment of renal epithelial cell cilia, resulting in polycystic kidneys (17, 18). In addition, YAP/TAZ inhibition by verteporfin ameliorates renal fibrosis after unilateral ureteral obstruction in adult mice (19).

We and others have lately reported that a canonical Hippo pathway finely regulates development, differentiation, and maintenance of integrity of several stromal cells, including blood and lymphatic endothelial cells and fibroblasts (20–29). YAP/TAZ are essential not only for sprouting angiogenesis but also for maturation of the blood–retinal barrier and blood– brain barrier (21, 23). YAP/TAZ act as negative regulators of Prox1 in developmental lymphangiogenesis and lymphatic differentiation and in maintaining lymphatic integrity in adults (27). Although they are the main drivers of fibroblast activation and fibrosis in lung (22, 30, 31), YAP/TAZ also regulate vascular endothelial growth factor-C in distinct sets of villi fibroblasts for lacteal integrity in intestine (29). Moreover, both YAP and TAZ are highly enriched in fibroblastic reticular cells (FRCs) and regulate their commitment and maturation in the lymph nodes (28). Hyperactivation of YAP/TAZ transforms FRCs into dedifferentiated myofibroblasts and leads to excessive lymph node fibrosis (28).

The structure and roles of FRCs in the lymph nodes are similar to those of MCs in the renal glomerulus. In addition, both MCs and FRCs are constantly exposed to dynamic and versatile mechanical and biochemical stimuli, which are conveyed and transduced into the cells through the Hippo pathway. Therefore, we hypothesized that the MC Hippo pathway would be critically involved in not only regulating MC integrity but also the pathogenesis of renal diseases, including DN. Here, we show that the Hippo pathway is an important regulator in maintaining MC integrity and that over-activation of YAP/TAZ is crucially involved in DN pathogenesis.

## Results

### YAP and TAZ proteins are highly enriched in glomerular platelet-derived growth factor receptor (PDGFR)β^+^ MCs of patients with DN and in Zucker diabetic fatty (ZDF) rats

To determine whether the Hippo pathway is involved in MCs of DN, we first examined protein levels of YAP and TAZ in paraffin-sectioned glomeruli obtained from five patients with DN and five unaffected portions of samples from patients with renal cell carcinoma (Table 1). Dual immunohistochemical staining for YAP or TAZ and PDGFRβ revealed that YAP and TAZ were highly enriched (42.7% and 18.8% in all PDGFRβ^+^ MCs) in the glomeruli from DN samples but rarely detected (6.8% and 0.0% in all PDGFRβ^+^ MCs) in the normal glomerulus tissues (Figure 1, A–D). To determine whether the diabetic condition in patients with DN is associated with increased levels or nuclear localization (activated) of YAP and TAZ in MCs, we analyzed the glomeruli of ZDF rats, which have a mutation in the leptin receptor gene resulting in insulin resistance and reduced glucose tolerance, and thus in DN (32, 33). As control, we used Zucker diabetic lean (ZDL) rats of the same age (22 weeks). Compared with ZDL rats, ZDF rats had 30% less body weight (BW) and 2.5-fold higher fasting blood glucose, along with hypercellular glomeruli with increased mesangial proteoglycan and collagen deposition (Figure 2A and Supplemental Figure 1). These manifestations suggested that diabetes mellitus and DN developed in the ZDF animals. Of note, compared with the PDGFRβ^+^ MCs of ZDL rats, those of the ZDF group had 12.0-fold and 4.2-fold increased nuclear-localized YAP and TAZ, respectively, along with 17.0-fold and 4.0-fold increased Ki67^+^ and EdU^+^, respectively. Moreover, glomeruli of ZDF rats had 2.1-fold higher collagen IV and 8.5-fold higher αSMA^+^ PDGFR β^+^ MCs (Figure 2, B–D). These findings imply that YAP/TAZ can be hyperactivated in the diabetic condition, which could lead to abnormal proliferation and de-differentiation of MCs into myofibroblasts.

**Figure 1.**
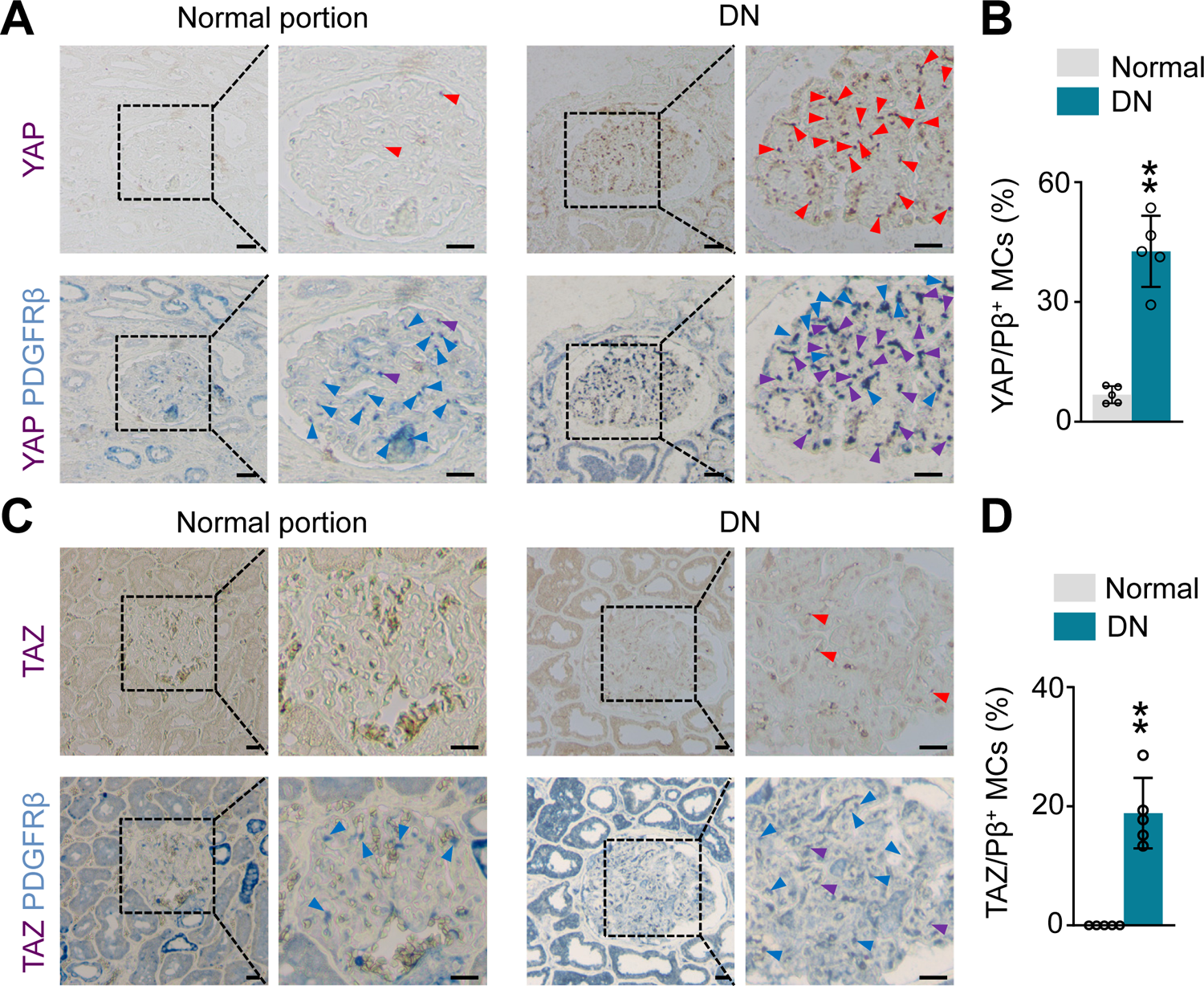
YAP^high/^TAZ^high^ PDGFRβ^+^ MCs are abundantly detected in the glomeruli of patients with diabetic nephropathy. (A) Representative images and comparisons of YAP^high^ PDGFRβ^+^ (YAP/ Pβ^+^) MCs (red arrowheads in upper panels and purple arrowheads in lower panels) in the glomeruli at the unaffected region of the patients of renal carcinoma (RCC) *versus* the patients with DN. Blue arrowheads indicate YAP^No^PDGFRβ^+^ MCs. (B) Representative images and comparison of TAZ^high^ PDGFRβ^+^ (TAZ/ Pβ^+^) MCs (red arrowheads in upper panels and purple arrowheads in lower panels) in the glomeruli at the normal region of the patients of RCC *versus* the patients with DN. Blue arrowheads indicate TAZ^No^PDGFRβ^+^ MCs. All scale bars, 20 μm. Each dot indicates a value from one patient and n = 5/group from three independent experiments. Vertical bars indicate mean ± SD. **P<0.01 *versus* normal glomeruli obtained from RCC patient by unpaired t-test.

**Figure 2.**
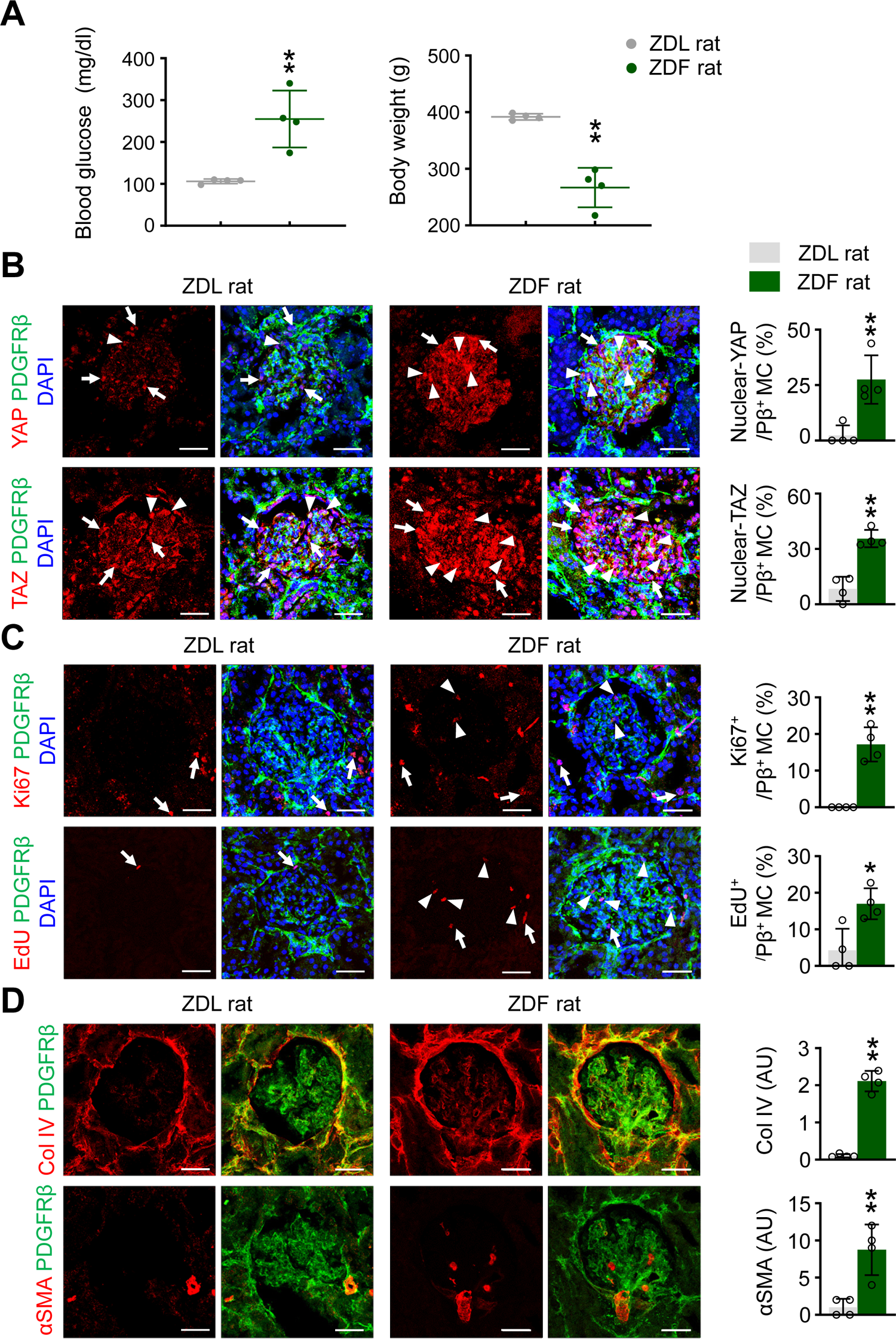
ZDF rats have high and activated YAP and TAZ in MCs and show hallmarks of DN in the renal glomeruli. (A) Comparisons of blood glucose (mg/dl) and body weight (g) of 22-week-old ZDL and ZDF rats. (B) Representative images and comparisons of YAP and TAZ and their subcellular localizations in the PDGFRβ+ MCs of glomeruli in 22-week-old ZDL and ZDF rats. Note that YAP and TAZ are highly localized in the nuclei (white arrowheads) of PDGFRβ+ MCs in glomeruli of ZDF rats. White arrows indicate nuclear YAP or TAZ in non-mesangial cells. (C) Representative images and comparisons of number of Ki67^+^ or EdU^+^ proliferating PDGFRβ+ MCs per glomeruli in 22-week-old ZDL and ZDF rats. White arrowheads indicate Ki67^+^ and EdU^+^ in mesangial cells. White arrows indicate Ki67^+^ and EdU^+^ non-mesangial cells. (D) Representative images and comparisons of collagen type IV (Col IV) and αSMA in mesangial cells of 22-week-old ZDL and ZDF rats. All scale bars, 20 μm. Each dot indicates a value from one rat and n = 4 rats/group from two independent experiments. All vertical bars indicate mean ± SD. *P<0.05 and **P <0.01 *versus* ZDL rats by unpaired t-test.

**Table 1.**
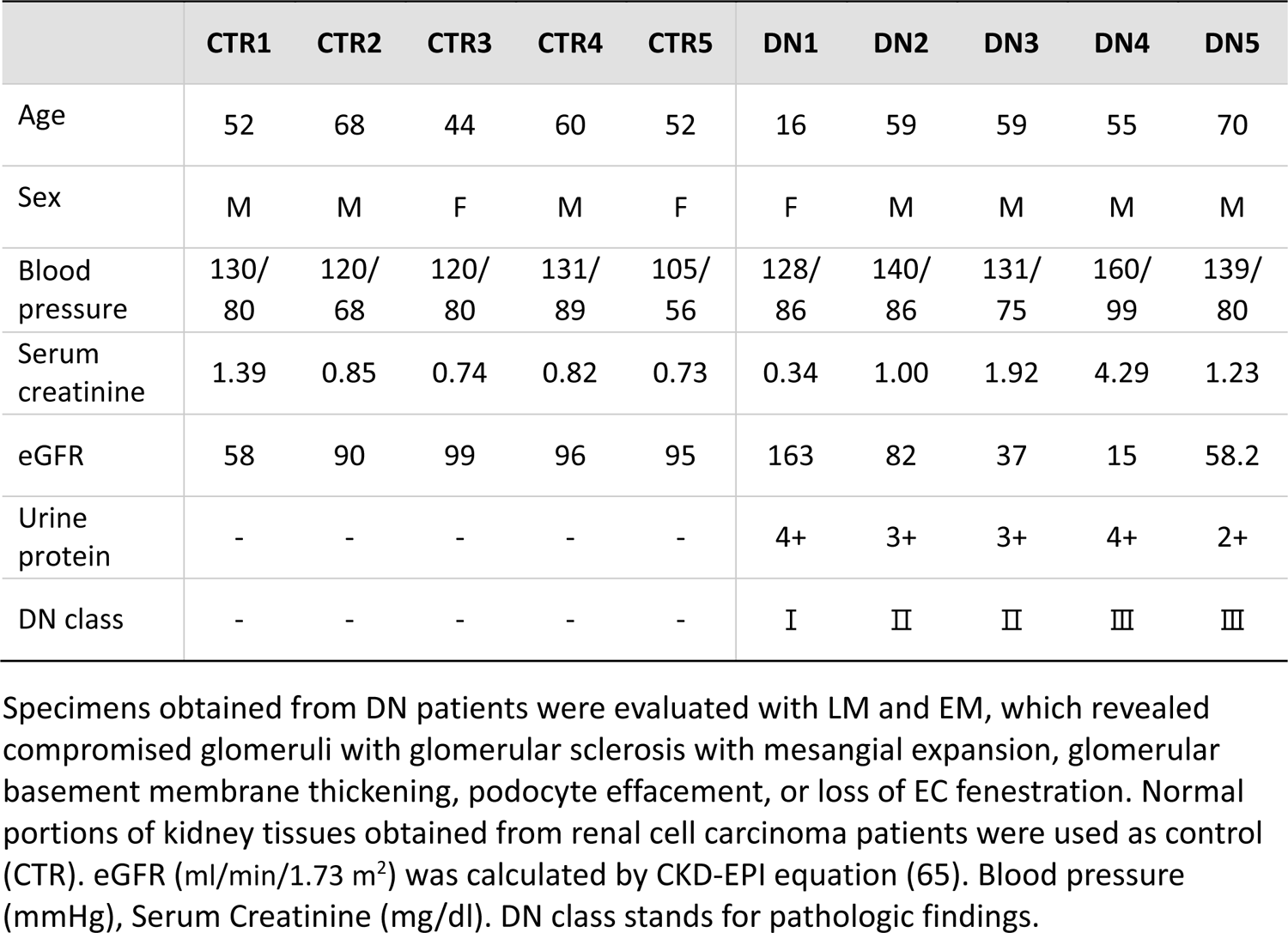
Comparisons for clinical characteristics of subjects.

To delineate whether high blood glucose regulates the Hippo pathway in MCs, we exposed immortalized mouse MCs, the SV40-MES13 cell line (34), to various glucose concentrations. Upon glucose exposure, phosphorylated MST1, LATS1, and YAP were reduced in a concentration-dependent manner (Figure 3, A and B). Moreover, high glucose (50 mM) caused 3.2-fold increased nuclear localization of YAP compared with a normal glucose concentration (5.5 mM) (Figure 3, C and D). Thus, high glucose exposure directly induced activation of YAP/TAZ through the canonical Hippo pathway in cultured MCs.

**Figure 3.**
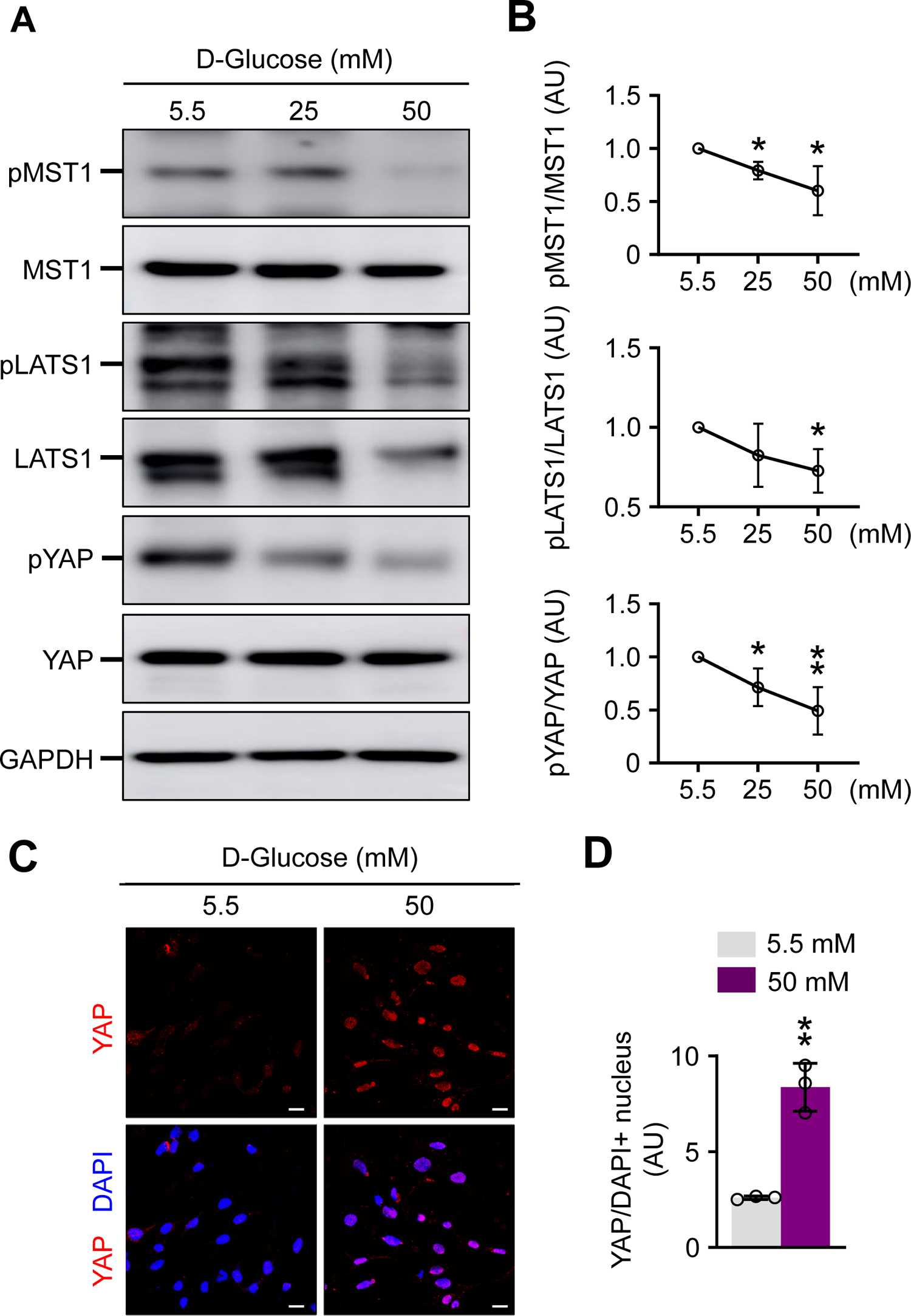
YAP/TAZ are activated by high D-glucose in a canonical Hippo pathway in cultured MCs. (A) Represeantative immunoblotting analysis showing reduced pMST1, pLATS1 and pYAP in a D-glucose concentration dependent manner in cultured mouse MC lines, SV40-MES13 for 24 h. (B) Relative intensities of pMST/MST, pLATS1/LATS1 and pYAP/YAP. Dots and bars indicate mean ± SD and n = 3/group from three independent experiments. *P<0.05 and **P<0.01 *versus* 5.5 mM D-glucose by unpaired t-test. (**C** and **D**) Representative images and comparisons of nuclear YAP in cultured SV40-MES13 cells, which are exposed to normal (5,5 mM) and high (50 mM) D-glucose medium for 24 h. Scale bars, 20 μm. Each dot indicates a value from one cell culture experiment and n = 3/group from two independent experiments. Vertical bars indicate mean ± SD. **P<0.01 *versus* 5.5 mM D-glucose by unpaired t-test.

### YAP/TAZ hyperactivation in PDGFRβ^+^ MCs promotes MC proliferation and glomerular fibrosis

The above findings led us to assess the roles of activated YAP/TAZ in the MCs. To target MCs for inducible YAP/TAZ activation *in vivo*, we used the *Pdgfrb*-Cre-ER^T2^ mouse (35) and *Lats1^fl/fl^/Lats2^fl/fl^* mouse (36, 37). First, to ensure that the MCs expressed PDGFRβ, we generated tdTomato^rPβ^ reporter mice by crossing *Pdgfrb*-Cre-ER^T2^ mice with *Rosa26-* tdTomato reporter mice, and injected the animals with tamoxifen for 3 days to activate PDGFRβ expression in the targeted kidney cells (Supplemental Figure 2A). We found that the PDGFRβ^+^ cells were NG2^+^ MCs in the glomeruli (Supplemental Figure 2B) (3). In comparison, the positive cells were pericytes and fibroblasts in peritubular capillary, NG2^+^ pericytes in the vasa recta bundle, and NG2^+^/α-SMA^+^ SMCs in kidney arteriole (Supplemental Figure 2B).

Taking advantage of the *Pdgfrb*-Cre-ER^T2^ mouse, we genetically targeted MCs. To gain insight into the role of YAP/TAZ activation in PDGFRβ^+^ MCs, we generated *Lats1/2^i^*^ΔPβ^ mice by crossing *Pdgfrb*-Cre-ER^T2^ mice with *Lats1^fl/fl^/Lats2^fl/fl^* mice (Figure 4A). Cre-ER^T2^ positive but flox/flox-negative mice among the littermates were used as control mice. We found that YAP/TAZ were rarely (<2%) detectable in PDGFRβ^+^ MCs from control mice but were highly and predominantly present (35%–65%) in the nuclei of PDGFRβ^+^ MCs from *Lats1/2*^iΔPβ^ mice (Figure 4B). The numbers of Ki67^+^ and EdU^+^ proliferating MCs increased by 20.1-fold and 19.0-fold in *Lats1/2*^iΔPβ^ mice compared with control mice values (Figure 4C), indicating that PDGFRβ^+^ MCs were actively proliferating in *Lats1/2*^iΔPβ^ mice. Moreover, collagen type I increased by 1.6-fold in *Lats1/2*^iΔPβ^ mice (Figure 4D), indicating excessive ECM deposition in the mesangium of these mice compared with control mice. Furthermore, αSMA^+^PDGFR β^+^ MCs were increased in the glomeruli of *Lats1/2*^iΔPβ^ mice by 3.5-fold (Figure 4D), implying that the proliferative PDGFRβ^+^ MCs were de-differentiated into myofibroblasts (38, 39). Taken together, these findings indicate that YAP/TAZ hyperactivation in PDGFRβ^+^ MCs induces their abnormal proliferation and de-differentiation into myofibroblasts, along with glomerular fibrosis, the common pathologic features of DN (11, 13).

**Figure 4.**
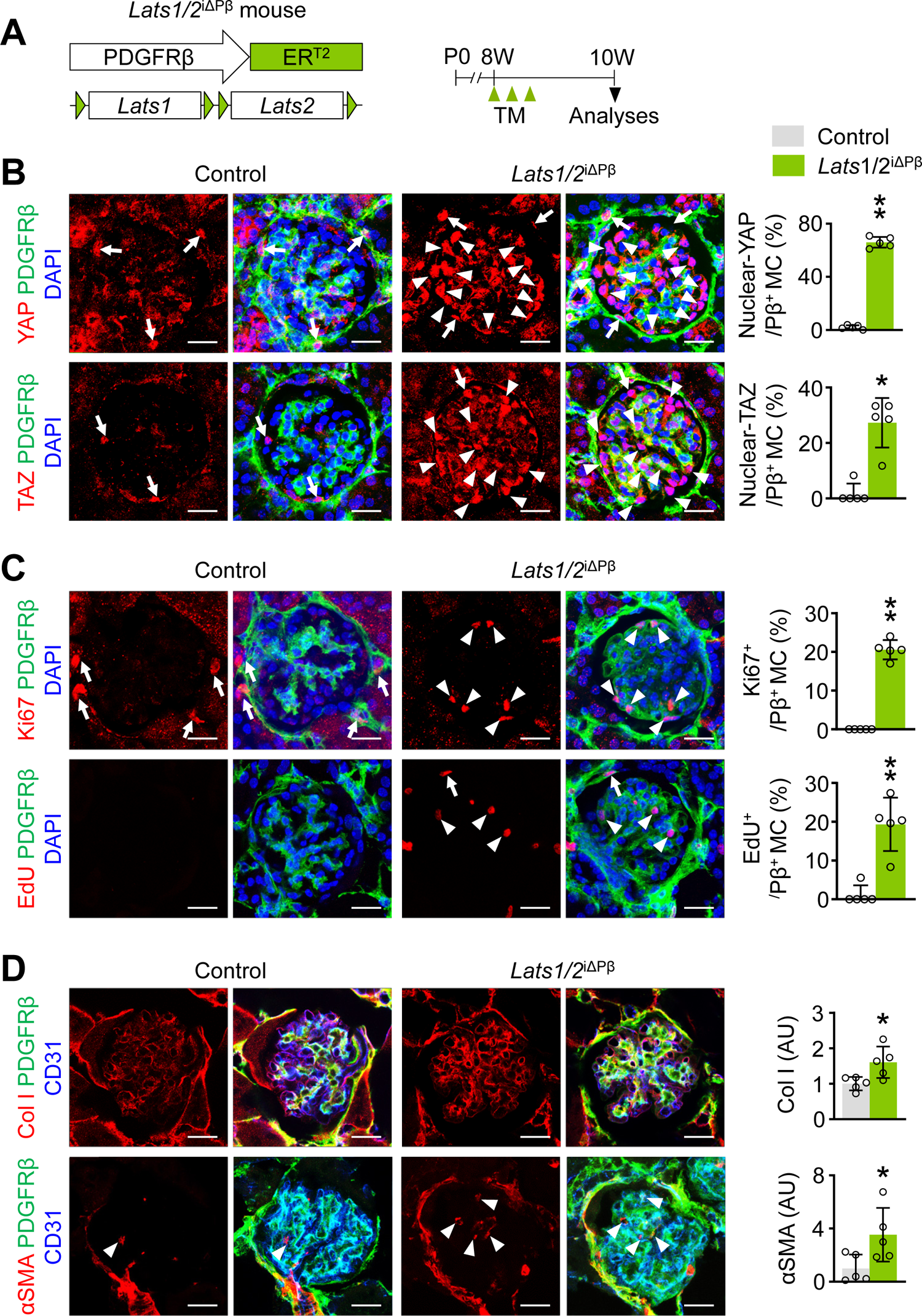
YAP/TAZ hyperactivation drives proliferation and fibrosis of MCs. (A) Diagram depicting the PDGFRβ^+^ cell-specific depletion of *Lats1/2* in *Lats1/2*^iΔPβ^ mice by tamoxifen (TM) administration from 8-week-old and their analyses at 10-week-old. (B) Representative images and comparisons of YAP and TAZ and their subcellular localizations in the PDGFRβ^+^ MCs of glomeruli in control and *Lats1/2*^iΔPβ^ mice. Note that YAP and TAZ are highly localized in the nuclei (white arrowheads) of PDGFRβ^+^ MCs in glomeruli of *Lats1/2*^iΔPβ^ mice. White arrows indicate nuclear-localized YAP or TAZ in non-mesangial cells. (C) Representative images and comparisons of number of Ki67^+^ or EdU^+^ proliferating PDGFRβ^+^ MCs per glomeruli in control and in *Lats1/2*^iΔPβ^ mice. White arrowheads indicate Ki67^+^ or EdU^+^ in mesangial cells, while white arrows indicate Ki67^+^ or EdU^+^ in non-mesangial cells. (D) Representative images and comparisons of collagen type I (Col I) and αSMA in mesangial cells of control and *Lats1/2*^iΔPβ^ mice. White arrowheads indicate αSMA^+^PDGFRβ^+^ MCs. All scale bars, 20 μm. Each dot indicates a value from one mouse and n = 5 mice/group from three independent experiments. All vertical bars indicate mean ± SD. *P<0.05 and **P<0.01 *versus* control by unpaired t-test.

### YAP/TAZ hyperactivation in PDGFRβ^+^ MCs leads to pathologic changes in glomerular ECs

To delineate the impact on the surrounding glomerular capillary bed of the mesangial alterations induced by YAP/TAZ hyperactivation, we performed immunofluorescence staining (IFS) on the glomerular ECs of control and *Lats1/2*^iΔPβ^ mice. In *Lats1/2*^iΔPβ^ mice compared with control mice, angiopoietin-2, a marker of pathologic activation of ECs (40), was 2.0-fold higher, whereas Tie2 and VE-cadherin were 27% and 32% lower, respectively (Figure 5B). In addition, claudin-5 was 23% lower, but ICAM-2 was 1.3-fold higher in *Lats1/2*^iΔPβ^ mice compared with control mice (Figure 5B). Nevertheless, we detected no notable differences between *Lats1/2*^iΔPβ^ and control mice in levels or distribution patterns of podoplanin, a surrogate marker of podocytes in glomeruli (Figure 5C). However, GLUT1, a hypoxic marker (41), was 2.0-fold higher in *Lats1/2*^iΔPβ^ mice compared with control mice (Figure 5C). These findings indicate that pathologic alterations in MCs lead to secondary pathologic changes to the surrounding capillary bed but not to podocytes in the renal glomeruli of *Lats1/2*^iΔPβ^ mice.

**Figure 5.**
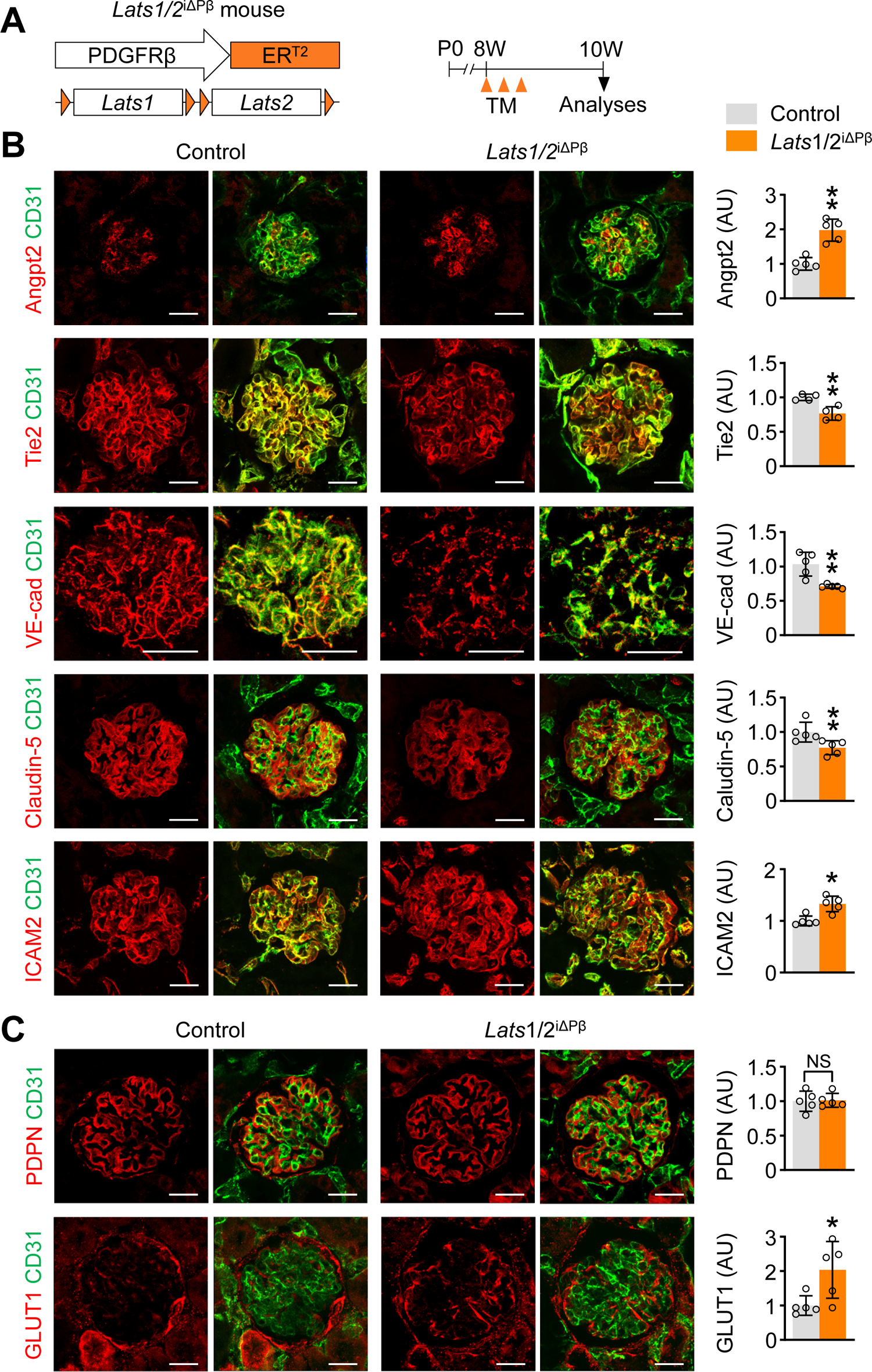
YAP/TAZ hyperactivation in PDGFRβ^+^ MCs impairs glomerular filtration barrier. (A) Diagram depicting the PDGFRβ^+^ cell-specific depletion of *Lats1/2* in *Lats1/2*^iΔPβ^ mice by TM administration from 8-week-old and their analyses at 10-week-old. (B) Representative images and comparisons of angiopoietin-2 (Angpt2), Tie2, VE-cadherin (VE-cad), claudin-5 and ICAM2 in glomeruli of control and *Lats1/2* ^iΔPβ^ mice. (C) Representative images and comparisons of podoplanin (PDPN) and GLUT1 in glomeruli of control and *Lats1/2*^iΔPβ^ mice. All scale bars, 20 μm. Each dot indicates a value from one mouse and n = 5 mice/group from three independent experiments. All vertical bars indicate mean ± SD. *P<0.05 and **P<0.01 *versus* control mice by unpaired t-test. NS, non-significant.

### YAP/TAZ hyperactivation in PDGFRβ^+^ MCs impairs glomerular filtration and induces glomerular sclerosis mediated through a canonical Hippo pathway

Compared with control mice, BW and kidney weight of *Lats1/2*^iΔPβ^ mice were 25% and 43% lower, respectively (Figure 6, A and B). Serum levels of cystatin-C, a biomarker for estimating glomerular filtration rate, and urine albumin were 1.5-fold and 13-fold higher in *Lats1/2*^iΔPβ^ mice versus controls (Figure 6B), indicating impairment of the glomerular filtration barrier in the *Lats1/2*^iΔPβ^ mice. Consistent with these functional alterations, compared with control mice, *Lats1/2*^iΔPβ^ mice showed hypercellular glomeruli with increased mesangial proteoglycan and collagen deposition (Figure 6C). As a result, *Lats1/2*^iΔPβ^ mice had a higher glomerular sclerosis index (Figure 6C). At the ultrastructural level, control mice showed a well-structured glomerular filtration barrier, but *Lats1/2*^iΔPβ^ mice exhibited swollen ECs (endotheliosis) and focal effacement of podocyte foot processes in the glomeruli (Figure 6D). Thus, the abnormal glomerular phenotypes and functional impairments of DN can be recapitulated by YAP/TAZ hyperactivation of PDGFRβ^+^ MCs in these mice.

**Figure 6.**
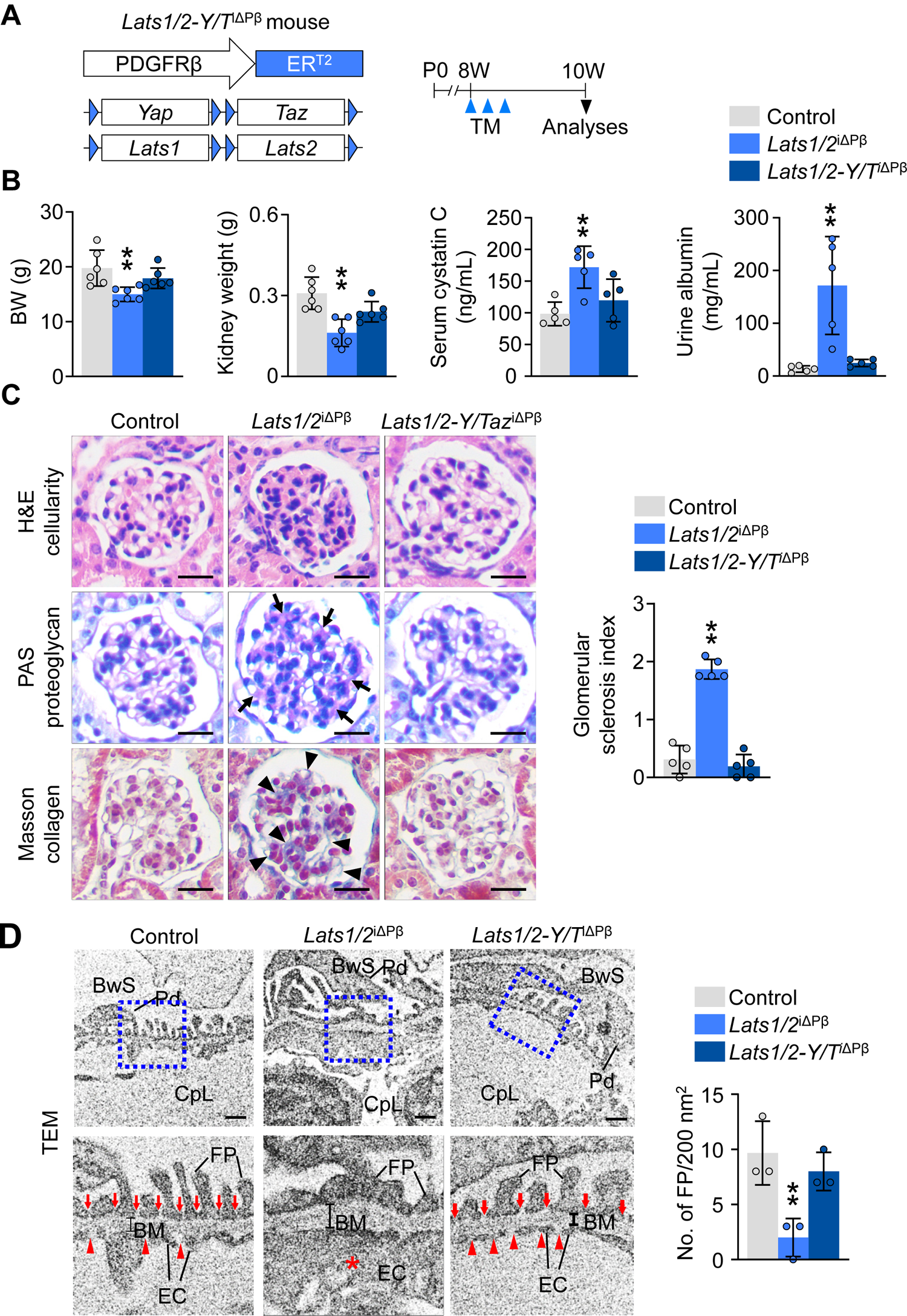
Co-deletion of *Yap/Taz* in MCs of *Lats1/2* ^iΔPβ^ mice rescues the glomerular pathologic phenotypes. (A) Diagram depicting the generation of *Lats1/2-Yap/Taz*^iΔPβ^ (*Lats1/2-Y/T*^iΔPβ^) mouse and PDGFRβ^+^ cell-specific depletion of *Lats1/2-Yap/Taz* by TM administration in 8-week-old mice and analyses at 2 weeks later. (B) Comparisons of body weight (BW), kidney weight, and concentrations of serum cystatin-C and spot urine albumin among control, *Lats1/2*^iΔPβ^ and *Lats1/2-Y/T*^iΔPβ^ mice. n = 6 mice/group from two independent experiments. (C) Representative images of H&E-stained cellularity, PAS+ proteoglycan (black arrows) and Masson’s Trichrome-stained collagen (black arrowheads) in the glomeruli and comparisons of glomerular sclerosis index among control, *Lats1/2*^iΔPβ^, and *Lats1/2-Y/T*^iΔPβ^ mice. Scale bars, 20 μm. n = 5 mice/group from three independent experiments. (D) Representative transmission electron micrographs (TEMs) of glomeruli and comparisons of numbers of foot process (FP) in control, *Lats1/2*^iΔPβ^ and *Lats1/2-Y/T*^iΔPβ^ mice. Each blue-lined box is magnified in below for visualization of detailed ultrastructure of glomerular filtration barrier. Note that silt diaphragms (red arrows) and endothelial fenestrations (red arrowheads) shown in control mice are disappeared in *Lats1/2*^iΔPβ^ mice, which leads to fusion and effacement of FP and endothelial cell swelling (endotheliosis, red asterisk). Silt diaphragms and endothelial fenestrations are restored in *Lats1/2-Y/T*^iΔPβ^ mice. n = 3 mice/group from three independent experiments. BwS, Bowman’s space; Pd, podocyte; CpL, capillary lumen; FP, foot process of podocyte; BM, basement membrane; EC, endothelial cell. Scale bars, 500 nm. Each dot indicates a value from one mouse. All vertical bars indicate mean ± SD. *P<0.05 and **P<0.01 *versus c*ontrol mice by one-way ANOVA followed by Dunnett’s multiple comparison test.

To delineate whether the *Lats1/2* deletion–induced glomerular pathologic alterations are directly mediated through YAP/TAZ hyperactivation in MCs, we generated quadruple *Last1/2*-*YAP/TAZ*^iΔPβ^ (*Last1/2*-*Y/T*^iΔPβ)^ mice by crossing *Pdgfrb*-Cre-ERT2 animals with *Last1^fl/fl^/Last2^fl/fl^* mice and *YAP*^fl/fl^/*TAZ*^fl/fl^ mice (42, 43), and compared their glomerular phenotypes (Figure 6A). Functional parameters including BW, kidney weight, serum cystatin-C, and urine albumin of *Last1/2*-*Y/T*^iΔPβ^ mice were comparable to those of control mice (Figure 6B). Furthermore, structural parameters such as glomerular cellularity, mesangial proteoglycan and collagen deposition, glomerular sclerosis, and the ultrastructural level of the glomerular filtration barrier of *Last1/2*-*Y/T*^iΔPβ^ mice were also similar to those of control mice (Figure 6, C–D). Thus, YAP/TAZ hyperactivation leads to alterations in PDGFRβ^+^ MCs through the canonical Hippo pathway.

### N-Myc signaling is upregulated in PDGFRβ^+^ MCs of *Lats1/2* ^iΔPβ^ mice

These findings led us to investigate how YAP/TAZ hyperactivation in PDGFRβ^+^ MCs causes glomerular pathologic alterations. To address this question, we used RiboTag RNA sequencing and performed transcriptomic analysis of gene expression profiles in isolated MCs from control and *Lats1/2*^iΔPβ^ mice. To obtain efficient and selective isolation of mRNA from PDGFRβ^+^ MCs, we generated RiboTag-*Lats1/2*^ΔPβ^ (RT-*Lats1/2*^ΔPβ^) and RiboTag^ΔPβ^ (control) mice by crossing RiboTag animals (44, 45) with *Lats1/2*^iΔPβ^ and wild-type mice (Figure 7A). We purified ribosome-associated actively transcribed mRNA in MCs by enzymatic digestion of renal cortical tissues, followed by isolation of glomeruli with magnetic beads (Figure 7B, Supplemental Figure 3).

**Figure 7.**
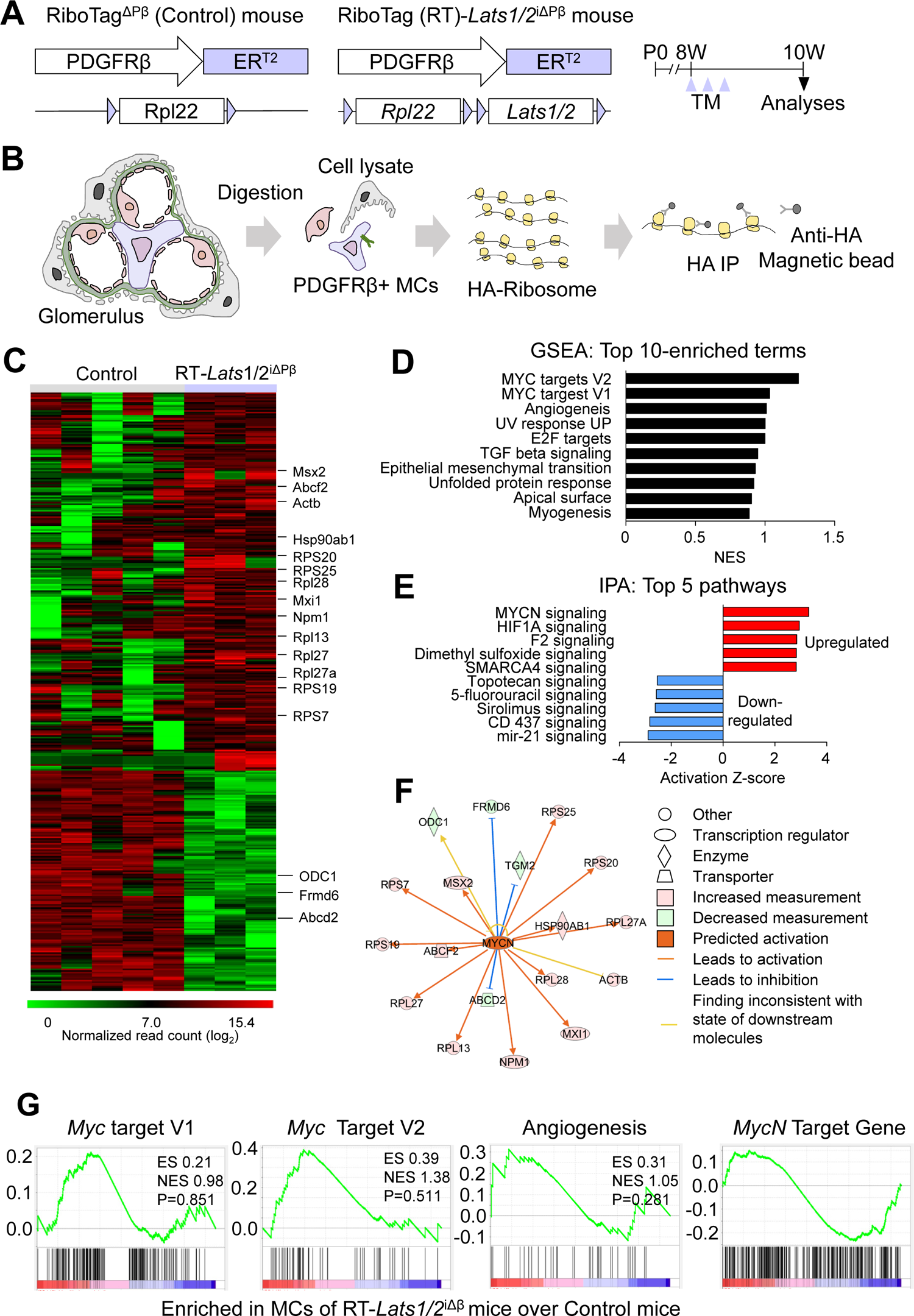
*MycN* signaling pathway is a downstream regulator of activated YAP/TAZ in MCs. (A) Diagram depicting generation of RiboTag^ΔPβ^ and RiboTag-*Lats1/2*^iΔPβ^ (inducible and specific tagging of ribosome in PDGFRβ^+^ cells) mice. TM administration from 8-week-old and their analyses at 10-week-old. (B) Diagram depicting a specific isolation of RiboTag mRNA from isolated PDGFRβ^+^ MCs of RiboTag^ΔPβ^ (Control) and RiboTag-*Lats1/2*^iΔPβ^ (RT-*Lats1/2*^iΔPβ^) mice. Inducible recombination of RiboTag allele lead to expression of hemagglutinin (HA) - tagged ribosomal protein specifically in PDGFR-β^+^ MCs. Ribosome bound transcripts are immunoprecipitated from homogenized glomerular tissues with anti-HA antibody-coupled magnetic beads. Extracted mRNAs are analyzed by RNA-Seq analysis. 3-5 mice/group were used. (C) Heatmap and hierarchical clustering of differentially expressed genes of RNA-Seq data in isolated MCs from Control (n=5) and RT-*Lats1/2*^iΔPβ^ (n=3) mice. Most highly expressed genes are indicated within each cluster pattern (right). (D) GSEA of the top 10 upregulated terms in the hallmark gene set from MCs of RT-*Lats1/2*^iΔPβ^ mice compared to MCs of Control mice. MYC target genes were highly enriched in RT-*Lats1/2*^iΔPβ^ mice. NES, normalized enrichment score. (E) Canonical IPA-annotated pathways listed in absolute IPA activation Z-score (P<0.05) to identify potential activation or inhibition of indicated signaling pathways in isolated MCs from RT-*Lats1/2*^iΔPβ^ compared with those of control mice. IPA z-score scale predicts activation or inhibition states of respective signaling pathways in MCs of RT-*Lats1/2*^iΔPβ^ mice. Top 5 upregulated or downregulated pathways are listed. (F) IPA network analysis indicates annotated interactions among the genes related to N-MYC, which are enriched in MCs of RT-*Lats1/2*^iΔPβ^ mice compared with those of Control mice. Up-(pink) and down-(green) regulated genes are indicated. The lines between genes represent known interactions, with colors representing activation and inhibition. Nodes are displayed using various shapes that represent the functional class of the gene product. (G) GSEA of indicated each target gene RT-*Lats1/2*^iΔPβ^ mice showed enriched *MYC* target gene set, genes related to angiogenesis, and *McyN* target gene set

Heat map analysis showed that genes related to cell proliferation and protein synthesis, including *Msx2*, *Abcf2*, *Hsp90ab1*, *Npm1*, and *Actb*, were 2.2 to 3.4-fold upregulated in RT-*Lats1/2*^ΔPβ^ mice compared with control mice (Figure 7C). Gene set enrichment analysis (GSEA) in PDGFRβ^+^ MCs of RT-*Lats1/2*^ΔPβ^ compared with control mice revealed increases in expression of genes related to *MYC* targets, angiogenesis, transforming growth factor-β signaling, and epithelial–mesenchymal transition (Figure 7, D and G). Of special note, Ingenuity Pathway Analysis (IPA) of differentially expressed genes revealed that the *MycN* signaling pathway was by far the most activated in PDGFRβ^+^ MCs of RT-*Lats1/2*^ΔPβ^ mice (Figure 7E). Moreover, IPA analysis revealed that molecules known to interact with N-MYC were up- and down-regulated, whereas GSEA analysis showed that *MycN* target genes were enriched in RT-*Lats1/2*^ΔPβ^ mice compared with control mice (Figure 7, F and G).

### YAP/TAZ induce proliferation and fibrosis through alteration of N-Myc stability

Because activation of N-MYC is related to proliferation of neural stem cells and fibroblasts and to pulmonary fibrosis (46–48), we hypothesized that N-Myc is a main target of the YAP/TAZ-driven proliferation of MCs and glomerular fibrosis. To test this hypothesis, we checked N-Myc in the PDGFRβ^+^ MCs of control and *Lats1/2* ^iΔPβ^ mice. Protein levels of N-Myc in the nuclei of PDGFRβ^+^ MCs were 5.7-fold higher in *Lats1/2*^iΔPβ^ mice compared with control mice (Figure 8, A and C). Nonetheless, nuclear TEAD4 levels in the PDGFRβ^+^ MCs were not different between the two groups (Figure 8, B and C).

**Figure 8.**
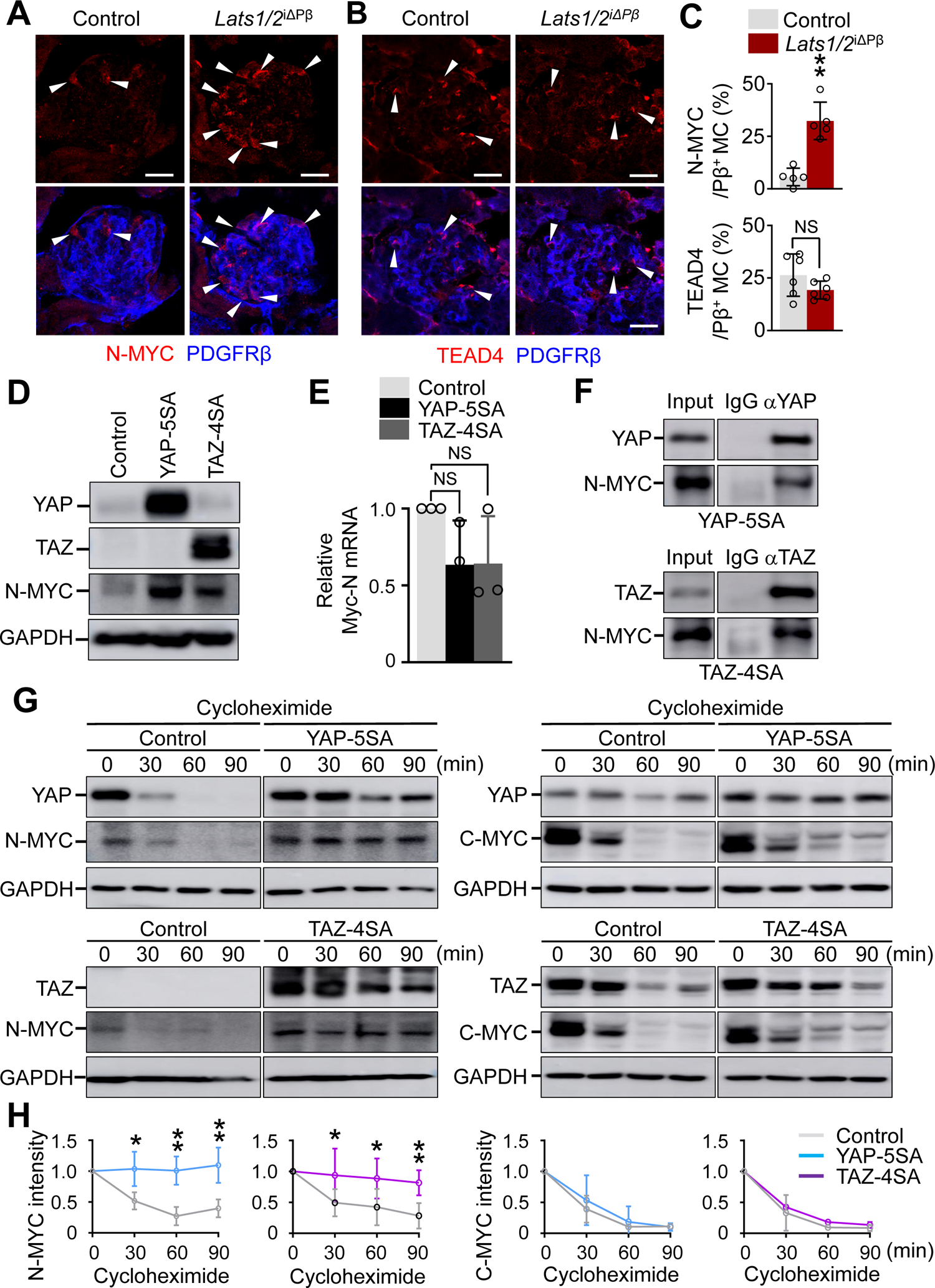
N-MYC is upregulated by YAP/TAZ through delaying protein degradation. (**A-C**) Representative images of N-MYC and TEAD4 and their subcellular localizations (white arrowheads) in PDGFRβ^+^ MCs of the glomeruli of Control and *Lats1/2*^iΔPβ^ mice. Scale bars, 20 μm. Each dot (for N-MYC) and two dots (for TEAD4) indicates values from one mouse and n=3-5 mice/group from 2-3 independent experiments. Vertical bars indicate mean ± SD. **P<0.01 *versus* control by unpaired t-test. NS, non-significant. (D) Immunoblot analysis of indicated proteins in cultured C3H/10T1/2 fibroblasts transfected with adenovirus-empty vector (Control), adenovirus-YAP5SA (AdeYAP-5SA) or adenovirus-TAZ4SA (AdeTAZ-4SA). n = 4/group from three independent experiments. (E) qRT-PCR analysis of *MycN* mRNA in cultured C3H/10T1/2 fibroblasts transfected with Control, AdeYAP-5SA or AdeTAZ-4SA. Each dot indicates a value from one qRT-PCR data and n = 3/group from three independent experiments. Vertical bars indicate mean ± SD. Non-significant (NS) *versus* Control by one-way ANOVA with Dunnett’s multiple comparisons test. (F) Immunoblotting of YAP, TAZ and N-MYC after immunoprecipitation with anti-IgG, anti-YAP or anti-TAZ antibody in cultured C3H/10T1/2 fibroblasts transfected with Control, AdeYAP-5SA or AdeTAZ-4SA. n = 3/group from three independent experiments. (**G** and **H)** Immunoblotting and comparisons for changes of YAP, TAZ, N-MYC and C-MYC at 0, 30, 60 and 90 min after treatment of cycloheximide (25 μg/ml) in cultured C3H/10T1/2 fibroblasts transfected with Control, AdeYAP-5SA or AdeTAZ-4SA. n = 3/group from three independent experiments. Dots and bars indicate mean ± SD. *P<0.05 and **P<0.01 *versus* control by unpaired t-test.

Hyperactivation of either YAP or TAZ, using transfection of adenoviruses encoding their constitutive active form (AdeYAP-5SA or AdeTAZ-4SA) (49) into C3H/10T1/2 fibroblasts, increased N-Myc protein, although neither one significantly changed mRNA level of *MycN* (Figure 8, D and E). Moreover, immunoprecipitation analysis with anti-YAP antibody and anti-TAZ antibody in the lysates of YAP-5SA-overexpressed or TAZ-4SA-overexpressed C3H/10T1/2 fibroblasts revealed that N-Myc was physically bound to YAP and TAZ (Figure 8F).

To investigate whether the protein stability of N-Myc is mainly regulated by YAP or TAZ after their interaction, we measured protein levels of N-Myc in cells with a high level of either protein after cycloheximide treatment. The half-life of N-Myc was longer with higher YAP or TAZ levels, indicating that YAP/TAZ stabilize N-Myc protein levels by reducing N-Myc degradation after forming the complex (Figure 8, G and H). In contrast, YAP or TAZ did not modify the half-life of C-Myc (Figure 8, G and H), which is consistent with previous findings (50).

### N-Myc promotes proliferation and ECM deposition of fibroblasts

To examine the direct roles of N-Myc in fibroblasts, we transduced C3H/10T1/2 fibroblasts with an empty retrovirus MSCV vector (Control) or retrovirus MSCV vector encoding *MycN* (MSCV-*MycN*). High levels of N-Myc were validated by immunoblotting analysis (Figure 9A). To analyze the proliferation rate, we conducted cytometric analysis of CFSE dilution. Compared with Control-transduced cells, C3H/10T1/2 fibroblasts transduced with MSCV-*MycN* showed 85% diminished mean fluorescence intensity (Mean FI) (Figure 9B), indicating that the MSCV-*MycN*-transduced fibroblasts underwent accelerated cell cycles. Moreover, deposition of cytoplasmic collagen type I was enhanced 2.7-fold in MSCV*-MycN-*transduced C3H/10T1/2 fibroblasts compared with Control–transduced cells (Figure 9C). Collectively, these results suggest that YAP/TAZ induce proliferation of MCs and expansion of mesangium via enhanced stabilization of N-Myc (Figure 9D).

**Figure 9.**
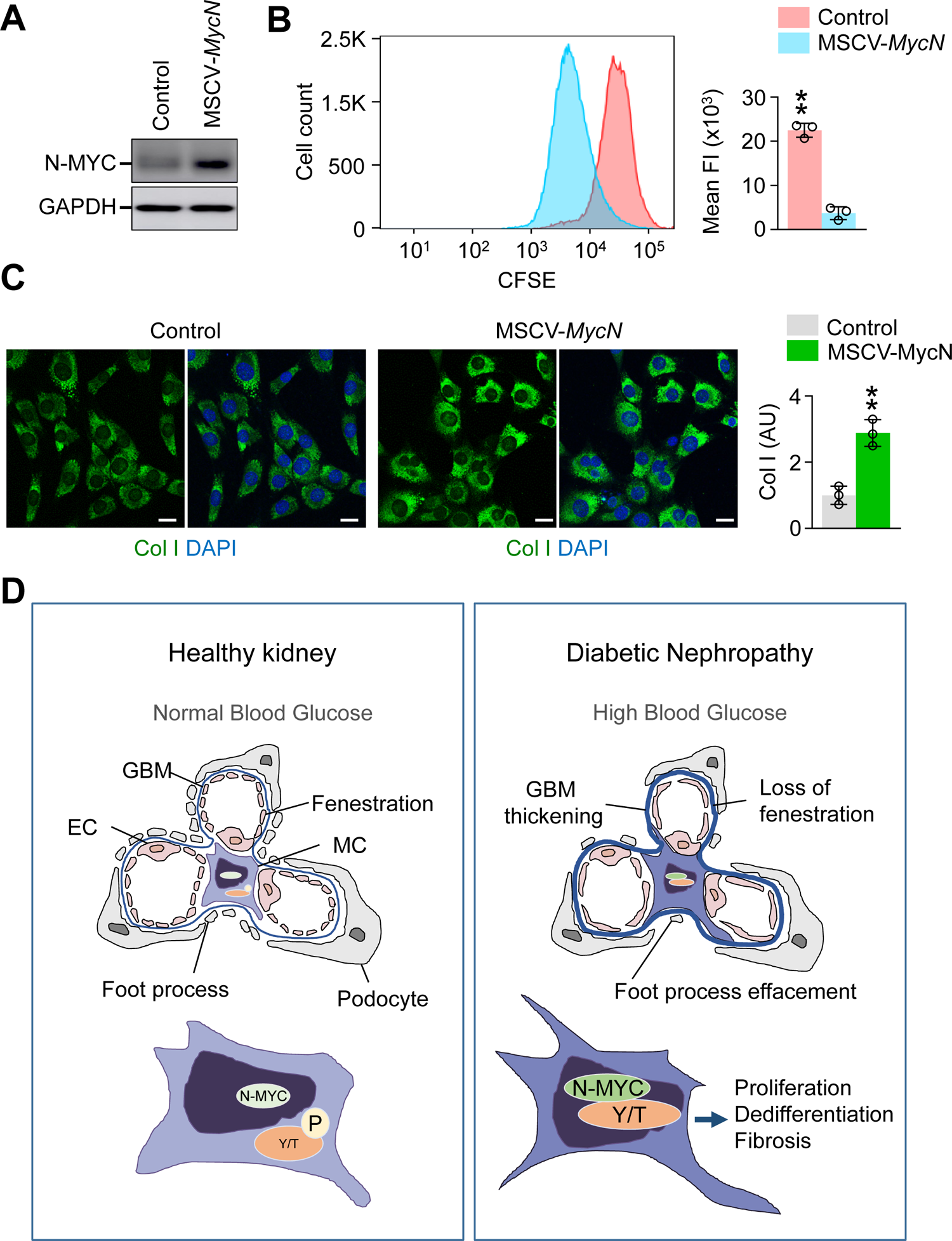
N-MYC promotes proliferation and collagen deposition of C3H/10T/1/2 fibroblasts. (**A** and **B**) Immunoblotting of N-MYC and CFSE dilution analysis in cultured C3H/10T1/2 fibroblasts transduced with MSCV-empty Vector (Control, pink) or MSCV-*MycN* (blue). Similar results were obtained from 3 independent blotting assays. CFSE fluorescence intensity (FI) histogram of each peck is shown. Note that MSCV-*MycN*-transduced cells show less FI compared with Control-transduced cells due to more proliferation. n = 3/group from three independent experiments. Vertical bars indicate mean ± SD. **P<0.01 versus Control by unpaired t-test. (C) Representative images and comparisons of collagen type I (Col I) level in C3H/10T1/2 fibroblasts transduced with Control vector or MSCV-*MycN.* Scale bar, 20 μm. n = 3/group from three independent experiments. Vertical bars indicate mean ± SD. *P<0.01 versus control by unpaired t-test. (D) Schematic diagram depicting how does the patient with diabetes mellitus (DM) get DN. High blood glucose of DM promote nuclear localization (activates) of YAP/TAZ in MCs in a canonical Hippo pathway, which leads to N-MYC activation through promoting its stability. Hyperactivation of N-MYC leads to proliferation, dedifferentiation and collagen overproduction in MCs, which eventually cause impairments of glomerular filtration barrier. GBM, glomerular basement membrane; EC, endothelial cell; MC, mesangial cell.

## Discussion

In this study, we found that YAP/TAZ proteins are highly enriched in MCs of patients with DN and in ZDF rats, and that high glucose directly induces activation of YAP/TAZ in a canonical Hippo pathway in cultured MCs. Accordingly, when we hyperactivated YAP/TAZ in MCs of genetically altered mice, we detected hallmarks of DN, such as excessive mesangial expansion and deposition of ECM, EC and podocyte impairments, glomerular sclerosis, albuminuria, and reduced glomerular filtration rate. Mechanistically, YAP/TAZ bind and stabilize N-Myc protein, aberrantly enhancing its transcriptional activity in MCs, and eventually leading to DN pathogenesis (Figure 9D).

Metabolites, metabolic pathways, and metabolic hormones play roles in regulating the Hippo-YAP signaling network, which integrates both local and systemic cues in a context-dependent manner (51). Glucose is a major source of cellular energy. Although YAP is usually phosphorylated and inactivated in a low glucose state (14), it is dephosphorylated and activated in a high glucose state in several cell types including ECs (52, 53). Consistent with these findings, we show that high glucose induces activation of YAP/TAZ in cultured MCs and in MCs from patients with DN and from ZDF rats. Moreover, high glucose increases O-GlcNAcylation of YAP and consequently enhances its expression, stability, and function, making it a potent oncogenic factor in tumorigenesis (54).

To delineate the roles of nuclear-enriched YAP/TAZ in MCs from patients with DN, we generated and analyzed outcomes in *Lats1/2^i^*^ΔPβ^ mice. This animal model faithfully recapitulates the glomerular impairment of DN (11, 13). Of note, the primary pathologic alterations in MCs induced by hyperactivation of YAP/TAZ seemed to cause secondary pathologic alterations in glomerular ECs, implying that MCs could be a primary and direct target in DN pathogenesis. Our genetic analysis revealed that YAP/TAZ hyperactivation in PDGFRβ^+^ MCs induces glomerular pathologic alterations mediated through a canonical Hippo pathway. Moreover, results of the transcriptomic, biochemical, and cellular analyses indicate that N-MYC could be the main driver in the YAP/TAZ hyperactivation-induced alterations of MCs.

N-MYC is one of the MYC family of oncogenes. These oncogenes encode basic helix-loop-helix zipper proteins and regulate expression of genes involved in cellular proliferation, growth, apoptosis, energy metabolism, and differentiation (55). C-MYC and N-MYC can replace each other in an appropriate context, but their functions are tissue-specific (56). C-MYC is highly expressed in most proliferating cells during development and continuously expressed in the dividing cells of adult tissues, whereas N-MYC shows restricted expression during neurogenesis and is downregulated in adult tissues (48, 55). Although C-MYC is a strong fibrosis factor (57, 58), the role of N-MYC in fibrosis has been unknown. In this study, we demonstrate that activated YAP/TAZ and N-MYC form a complex to enhance the stability of N-MYC, whereas activated YAP/TAZ do not affect the half-life of C-MYC.

Taken together, our findings indicate that activated YAP/TAZ induced by high glucose leads to sustained and high N-MYC activity, which eventually induces proliferation of MCs and glomerular fibrosis in patients with DN.

## Methods

### Human kidney tissues and immunohistochemistry (IHC)

Human kidney tissues were collected from the patients undertaken nephrectomy or renal biopsy due to renal cell carcinoma or DN, which was approved by the Institutional Review Board of Jeonbuk National University Hospital (CUH-2021-01-044-002). The tissues were paraffin-embedded, cut into 4 μm-thickness sections, and arrayed on a gelatin-coated slide. The sectioned slides were de-paraffinized, antigen-retrieved with a boiled 10 mM sodium citrate buffer (pH 6.0), incubated with hydrogen peroxidase for 40 min to block endogenous peroxidase, and incubated with 5% FBS in 0.2% Triton X-100 in PBS (PBST) for 1 h at room temperature (RT) for blocking of nonspecific binding of the antibodies. They were incubated with the primary anti-YAP antibody (rabbit, monoclonal, D8H1x, Cell signaling, 14074) or anti-TAZ antibody (rabbit, polyclonal, Sigma, HPA007415; dilution, 1:200 in blocking solution) for 48 h at 4°C. After wash with PBS, they were incubated with anti-rabbit-HRP conjugated antibody (R&D systems, HAF 008; dilution, 1:1,000 in blocking solution) for 1 h at RT. They were washed with PBS, and then incubated in DAB solution (DAKO, #K0679) for 1 min. The slides were washed with PBS and images were acquired using AxioZoom V16 fluorescence stereo zoom microscope (Carl Zeiss). To distinguish MCs in the glomeruli, a dual staining was performed by addition IHC of PDGFRβ in the primary stained slides for YAP or TAZ detection. The slides were rinsed with PBS, blocked with blocking solution for 1 h at RT, and incubated with biotin-conjugated anti-PDGFRβ antibody (Invitrogen, 13-1402-82; dilution, 1:200 in blocking solution) for 48 h at 4°C. After washing with PBS, they were incubated with an ABC-AP kit (Vector, AK-5000) for 1 h at RT. After washing, they were incubated with the blue substrate (Vector, SK-5300) for 30 min at RT. The slides were dehydrated and mounted with a permanent mounting solution (Sigma-Aldrich, 1.07960). Dual stained Images were acquired using AxioZoom V16 fluorescence stereo zoom microscope.

### Animals

Animal care and experimental procedures were performed under the approval from the Institutional Animal Care and Use Committee of KAIST (KA2018-49 and KA2021-121). All procedures and animal handlings were performed following the ethical guidelines for animal studies. *Pdgfrb*-Cre-ER^T2^ (35), *Lats1^fl/fl^/Lats2^fl/fl^* (36, 37), and *YAP^fl/fl^/TAZ^fl/fl^* (42, 43) mice were transferred, established, and bred in specific pathogen-free animal facilities at KAIST. Cre-ER^T2^ positive but flox/flox-negative mice among the littermates were defined as control mice for each experiment. R26-tdTomato mice and RiboTag mice were purchased from Jackson Laboratory (Jackson Lab, Bar Harbor, ME). *Pdgfrb*-Cre-ER^T2^ mice were bred into a background of R26-tdTomato and RiboTag mice. In order to induce Cre activity in the Cre-ER^T2^ mice, 2 mg of tamoxifen (Sigma), dissolved in corn oil (Sigma), was injected intraperitoneally (i.p.) at indicated time point for each experiment. All mice were maintained in the C57BL/6 background and fed with free access to a standard diet (PMI Lab Diet). They were anesthetized with i.p. injection of a combination of anesthetics (80 mg/kg ketamine and 12 mg/kg of xylazine) before sacrifice. ZDL rats and ZDF rats were purchased from Charles River Laboratories (Yokohama, Japan) at 8 weeks of age. The rats were properly housed, handled daily, and kept in a controlled standard temperature (22-23°C), humidity (60%), and light-dark cycles (12/12 h). Throughout the experiment, all rats were fed with free access to standard diet (Purina 5008 diet). Feeding was restricted one day before sacrifice. All rats were anesthetized with inhalation of isoflurane before sacrifice.

### Sample preparation and immunofluorescence staining (IFS)

After anesthesia, trans-cardial perfusion was performed into the mice and rats with PBS followed by 4% paraformaldehyde (PFA, Merck). The kidneys were harvested and the samples were post-fixed in 4% PFA at 4°C for 90 min. Samples were washed with PBS for several times and were subsequently dehydrated with 20% sucrose in PBS for overnight at 4°C. Samples were embedded in tissue freezing media (Lecia) and frozen blocks were cut in to 30 μm-thickness section. Samples were blocked with 5% donkey or goat serum in 0.3% PBST and incubated for overnight at 4°C with primary antibodies (dilution, 1:200 in blocking solution). After several washings with PBST, samples were incubated for 2 h at RT with secondary antibodies (dilution, 1:1,000 in blocking solution). For IFS of cultured SV40 MES13 cells and C3H/10T/1/2 cells, cells were plated in 4-well Nunc Lab Tek II chamber slides (Sigma-Aldrich) and fixed with 4% PFA for 10 min at RT. After several washings with PBS, samples were blocked with 5% goat or donkey in 0.2% PBS-T for 1 h at RT. Cells were incubated with primary antibodies (dilution, 1:200 in blocking solution) for overnight at 4°C. After several washes, cells were incubated with secondary antibodies (dilution, 1:1,000 in blocking solution) for 90 min at RT. The following primary antibodies were used for IFS: anti-YAP (rabbit, monoclonal, D8H1x, 14074, CST), anti-TAZ (rabbit, polyclonal, HPA 007415, Sigma), anti-PDGFRβ (rat, monoclonal, ab61066, Abcam), anti-PDGFRβ (goat, polyclonal, AF1042, R&D systems), anti-Ki67 (rabbit, monoclonal, SP6, ab16667, Abcam), anti-αSMA, fluorescein Isothiocyanate (FITC)-conjugated (mouse, monoclonal, S3777, Sigma-aldrich), anti-collagen type I (rabbit, polyclonal, ab6586, Abcam), anti-collagen type IV (rabbit, polyclonal, ab6586, Abcam), anti-CD31 (hamster, monoclonal, 2H8, Milipore), anti-CD31 (rat, polyclonal, 553370, BD), anti-angiopoietin-2 (human, monoclonal, clone 4H10) (59), anti-Tie2 (rabbit, monoclonal, 3A5, Santa Cruz, sc-293414), anti-VE-cadherin (goat, polyclonal, AF1002, R&D systems), anti-ICAM2 (rat, polyclonal, 553326, BD), anti-podoplanin (mouse, polyclonal, 127402, BioLegend), anti-claudin5 (rabbit, polyclonal, 34-1600, invitrogen), anti-GLUT1 (rabbit, polyclonal, 07-1401, Milipore), anti-N-MYC (rabbit, polyclonal, ab198912, abcam), anti-TEAD4 (mouse, monoclonal, 5H3, ab58310, abcam). Alexa fluor 488-, Alexa fluor 594-, Alexa fluor 647-conjugated anti-rabbit, anti-rat, anti-goat, anti-hamster, anti-human secondary antibodies were purchased from Jackson ImmunoResearch. Finally, the slides were mounted with a fluorescent mounting medium (DAKO) after several washings with PBST. Images were acquired using LSM 880 confocal microscope (Carl Zeiss).

### EdU incorporation assay for proliferating MCs

To detect proliferating MCs in glomeruli, 5 mg of 5-ethynyl-2′-deoxyuridine (EdU, A10044, Invitrogen) was dissolved in 1 ml of PBS as a stock solution. Then, 200 μl of the stock solution per mice and 1 ml of stock solution per rat were injected i.p. every other day for a week before analysis. Kidneys were isolated and processed as described above. EdU-incorporated cells were detected with the Click-iT EdU Alexa Fluor-488 Assay Kit (Invitrogen) according to the manufacturer’s protocol.

### Morphometric analyses

Morphometric analyses of the glomeruli were performed with images by photographic analysis using ImageJ software (http://rsb.info.nih.gov/ij) or ZEN 2012 software (Carl Zeiss).. Nuclear YAP or TAZ was counted by calculating the proportion of PDGFRβ^+^ MCs with nuclear YAP or TAZ from random five 100 μm^2^ fields of glomeruli and averaged Numbers of Ki-67^+^ and EdU^+^ proliferating MCs were measured from random five 100 μm^2^ fields of glomeruli and averaged. The relative expressions of indicated markers were calculated with dividing collagen type IV, αSMA, collagen type I, angiopoietin-2, Tie2, claudin5, ICAM2, podoplanin, and GLUT1 area by PDGFRβ^+^ or CD31^+^ area. For density measurements, VE-cadherin area was measured in random five 100 μm^2^ area and presented as a percentage of the total measured area.

### Cell culture and treatment

SV40-MES13 cells were purchased from ATCC (CRL 1927, ATCC) and cultured in 3:1 mixture of DMEM and Ham’s F-12 medium with 14 mM HEPES, containing 5% FBS. To mimic the diabetic environment, the media were supplemented with D-glucose (Sigma, G7021) at a final concentration of 5.5, 25, and 50 mM for 24 h. C3H/10T1/2 fibroblasts were purchased from ATCC (CCL-226, ATCC) and cultured in complete DMEM containing 10% FBS. For IFS of the cells, 1×10^4^ cells per well were plated on 4-well Nunc Lab-Tek II chamber slides (Sigma-Aldrich). The cultured SV40 MES13 cells were treated with 5.5 mM or 50 mM of D-glucose.

### Immunoblotting

For immunoblotting, cells were lysed with RIPA lysis buffer supplemented with protease and phosphatase inhibitors (Roche). Cell lysates were centrifuged at 15,000 rpm for 20 min at 4°C. Protein concentration of the supernatants was quantitated using the detergent-insensitive Pierce BCA protein assay kit (Thermo Fischer, 23227). 5X SDS loading buffer (LPS solution, CBS 002) was added to total protein lysate and samples were denatured at 95°C for 5 min. Aliquots of each protein lysate (20-30 μg) were subjected to SDS polyacrylamide gel electrophoresis. After electrophoresis, proteins were transferred to nitrocellulose membrane and blocked for 1 h with 2% BSA in 0.1% Tween 20 in TBS (TBST). After blocking, the membranes were incubated with following primary antibodies (dilution, 1:1,000 in blocking solution) at 4°C overnight; anti-MST1 (rabbit, polyclonal, Cell Signaling, 3682), anti-pMST1 (Thr183) (rabbit, monoclonal, E7U1D, Cell Signaling, 49332), anti-LATS1 (rabbit, monoclonal, C66B5, Cell Signaling, 3477), and anti-pLATS1 (Thr1079) (rabbit, monoclonal, D57D3, Cell Signaling, 8654), anti-YAP (rabbit monoclonal, D8H1x, Cell Signaling, 14074), anti-pYAP (Ser127) (rabbit, polyclonal, Cell signaling, 4911), anti-TAZ (rabbit polyclonal, Sigma, HAP-007415), anti-N-MYC (rabbit, polyclonal, Cell Signaling, 9405), anti-MYC (rabbit, polyclonal, Millipore, 06-340), anti-GAPDH (rabbit monoclonal, D16H11, Cell Signaling, 5174). After washings, membranes were incubated with anti-rabbit secondary peroxidase-coupled antibody (Cell signaling, 7074) for 90 min at RT. The signal intensity of targeted protein was detected using ECL western blot detection solution (Millipore, WBKLS0500).

### ELISA

Urine albumin and serum cystatin-C concentrations were measured by ELISA using commercial kits (Abcam, ab108792 for albumin; ab65340 for cystatin-C). Mouse urine and serum samples were freshly obtained from 8- to 10-week-old male control, *Lats1/2*^iΔPβ^, or *Lats1/2-YAP/TAZ*^iΔPβ^ mice, diluted 100-fold for albumin or 10-fold for cystatin-C, and measured according to the manufacturer’s instruction using a Spectra MAX340 plate reader (Molecular Devices).

### Measurement of glomerular sclerosis index

Kidney samples obtained from rats and mice were fixed in 2% PFA at 4°C for overnight. The fixed samples were embedded in paraffin, sectioned into 5 μm-thickness, and subjected to Hematoxylin and Eosin (H&E) staining, Periodic Acid-Schiff staining (PAS, Merck, 101646) and Masson’s trichrome staining (IHC world, IV-3006) according to manufacturer’s instructions, respectively. The glomeruli on PAS or Masson’s trichrome staining were evaluated “glomerular sclerosis” under 400x magnification using a semi-quantitative scoring from 0 to 4: Grade 0, normal appearance; Grade 1, involvement of up to 25% of the glomerulus; Grade 2, involvement of 25–50% of the glomerulus; Grade 3, involvement of 50–75% of the glomerulus; and Grade 4, involvement of 75–100% of the glomerulus. Glomerular sclerosis index (GSI, %) was calculated using the following formula (60), (1×n1/n+2×n2/n+3×n3/n+4×n4/n)×100, where nx is the glomeruli number in the rat kidney with a given score (x), n is the total glomeruli number.

### Electron microscopy

To capture ultrastructure electron microscopic (EM) images of glomeruli, kidney was sectioned after trans-cardial perfusion with 4% PFA and 0.25% glutaraldehyde in 0.1 M phosphate buffer (pH 7.4). Samples were then fixed overnight in 2.5% glutaraldehyde, post-fixed with 1% osmium tetroxide, and dehydrated with a series of increasing ethanol concentrations followed by resin embedding. 70-nm ultrathin sections were obtained using an ultra-microtome (UltraCut-UCT, Leica), which were then arrayed on copper grids. Samples were imaged with transmission EM (Tecnai G2 Spirit Twin, FEI) at 120 kV after staining with 2% uranyl acetate and lead citrate.

### mRNA isolation using RiboTag method

RiboTag mouse was used to isolate polysome-bound mRNAs of PDGFRβ^+^ MCs with a minor modification from previously described method (45). Upon tamoxifen treatment, RiboTag^ΔPβ^ mice tag hemagglutinin (HA) to the ribosome-associated actively transcribing mRNAs, specifically in PDGFRβ^+^ cells. Mouse kidneys were harvested and immediately snap frozen with liquid nitrogen. Then, polysome buffer (50 mM Tris, pH 7.5, 100 mM KCl, 12 mM MgCl2, 1% Nonidet P-40, 1 mM DTT, 200 U/ml RNasin, 1 mg/ml heparin, 100 μg/ml cycloheximide, and 1× protease inhibitor mixture) were added to each sample and homogenized using Precellys lysis kit (Bertin). Glomeruli were isolated with magnetic bead from whole lysates.

Cellular components of glomeruli were isolated using enzymatic digestion of kidney to excrete the mRNA, as previously described (61, 62). The finally purified product of digested kidney showed successful removal of non-glomerular structures from the primary digests of whole kidney and dissociation of cellular components (Supplemental Figure 3). After digestion of glomeruli, mRNA from PDGFRβ expressing cells of glomeruli was extracted and pulled down with antibody against HA tag. For IP against HA, anti-HA antibody-conjugated magnetic beads (MBL, M180-11) were added to the supernatant after centrifugation at 13,500g for 10 min at 4°C, and incubated on a rotating shaker at 4°C overnight. Beads were washed for 4 times with high-salt buffer (50 mM Tris, pH 7.5, 300 mM KCl, 12 mM MgCl2, 1% Nonidet P-40, 1 mM DTT, and 100 μg/ml cycloheximide) and resuspended in 350 μl of buffer RLT plus (Qiagen, 1053393) with β-mercaptoethanol. Total RNAs were extracted using Trizol RNA extraction kit (Invitrogen). The quality and quantity of the RNA samples were analyzed using Agilent 2100 Bioanalyzer with RNA 6000 pico kit (Agilent). Bulk-mRNA sequencing was carried out with these products.

### RiboTag RNA-sequencing

RiboTag RNA-sequencing of isolated PDGFRβ^+^ mesangial cells was performed by obtaining the alignment file. Briefly, reads were mapped using TopHat software tool. The alignment file was used to assemble transcripts, estimate their abundances, and detect differential expression of genes or isoforms using cufflinks. The normalized count values were processed based on Quantile normalization method using Genowiz and used for heatmaps and bioinformatic analysis. The IPA tool (QIAGEN) was used to further evaluate the data in the context of canonical signaling. The significance of the canonical signaling was tested by the Benjamini–Hochberg procedure (63), which adjusts the P value to correct for multiple comparisons, and their activation or inhibition was determined with reference to activation z-scores. For GSEA, gene set collections from the Molecular Signatures Database 4.0 (http://www.broadinstitute.org/gsea/msigdb/) were used.

### Adenoviral and retroviral transfections

C3H/10T1/2 cells that were serum starved for 12 h were transfected with ∼150 pfu/cell of AdeYAP-5SA or AdeTAZ-4SA (28, 49), and incubated them for 48 h prior to analysis. Mouse stem cell virus (MSCV)-based retroviruses encoding either mCherry alone (#52114; MSCV-mCherry) or co-express *MycN* and mCherry (#73571; MSCV-MycN-mCherry) were purchased from Addgene (64). Ten μg of retroviral constructs (pMSCV-*MycN*-mCherry, and pMSCV-mCherry) and two packaging plasmids (2 μg of VSV-G and 10 μg of pCMV-Gag-Pol) were co-transfected to HEK293T cells using lipofectamine LTX and PLUS reagent (lnvitrogen), and retroviral supernatant was collected 48 h after transfection. The supernatants were filtered through a 0.45-µm filter. The cultured C3H/10T1/2 cells were infected with 5 ml of retroviral supernatant for 48 h with 10 µg/ml Polybrene (Sigma-Aldrich).

### RNA extraction and quantitative RT-PCR

Total RNA was extracted from the C3H/10T1/2 cells using Trizol RNA extraction kit (Invitrogen). A total of 1 µg of extracted RNA was transcribed into cDNA using GoScript Reverse Transcription Kit (Promega). Quantitative real-time PCR was performed using FastStart SYBR Green Master mix (Roche) and QuantStudio3 (Applied Biosystems) with the indicated primers. *Gapdh* was used as a reference gene and the results were presented as relative expressions to control. Primer reaction specificity was confirmed by melting curve analysis. The real-time PCR data were analyzed with QuantStudio Software (Applied Biosystems). Results were calculated using the delta delta CT method. The primers were designed using Primer-BLAST: *MycN* (5’-ATTGGGCTACGGAGATGCTG-3’;5’-AGTCCTGAAGGATGACCGGA-3’), *Gapdh* (5’-TGATGGGTGTGAACCACGAG-3’;5’-GCCCTTCCACAATGCCAAAG-3’).

### Immunoprecipitation

The C3H/10T1/2 cells were harvested two days after the adenoviral infection and were lysed with NETN buffer [20 mM Tris-HCl (pH 7.4), 100 mM NaCl, 1 mM EDTA, 0.5% NP-40] with protease and phosphatase inhibitors (Roche). Cell lysates were centrifuged and the protein concentration of the supernatants were quantitated using BCA protein assay kit (Thermo Scientific, 23227). Cell extracts (800 μg) were incubated with one microliter of anti-YAP antibody (Santa Cruz Biotechnology, SCBT-101199) or anti-TAZ antibody (Cell Signaling, 4883) at 4°C overnight. Normal rabbit IgG (Cell Signaling, 2729) was used as a negative control. The immunoprecipitants were incubated with 50 µl of pre-washed protein A/G agarose bead (Thermo Fisher, 20421) for 4 h at 4°C. Immunoprecipitants with agarose bead were washed 3 times with NETN buffer, and boiled with Laemmli sample buffer at 95°C for 5 min. Aliquots of each protein lysate were subjected to SDS polyacrylamide gel electrophoresis. After electrophoresis, proteins were transferred to nitrocellulose membranes and blocked for 1 h with 2% BSA in TBST. Primary antibodies (dilution, 1:1,000 in blocking solution) were incubated overnight at 4°C. After several washings, membranes were incubated with True-Blot anti-Rabbit IgG HRP (ROCKLAND, Cat. 18-8816-31) for 1 h at RT. Target proteins were detected using ECL western blot detection solution (Millipore, WBKLS0500). Primary antibodies used for immunoblotting were as follows: anti-YAP (rabbit monoclonal, D8H1x, Cell Signaling, 14074), anti-TAZ (rabbit polyclonal, Sigma, HPA 007414), and anti-N-MYC (rabbit polyclonal, Cell Signaling, #9405).

### Cycloheximide chase assay

Cycloheximide (Sigma, c7698), a protein synthesis inhibitor, was used to analyze the stability of N-MYC and C-MYC. C3H/10T1/2 cells, transfected with each adenoviral vector (∼150 pfu/cell), were treated with cycloheximide (25 μg/ml) for 0, 30, 60, and 90 min. Proteins were extracted at the indicated time and immunoblot was performed to determine N-MYC and C-MYC level. The intensity of N-MYC and C-MYC was measured with ImagJ software (http://rsb.info.nih.gov/ij).

### Flow cytometric analysis of CFSE dilution

Two days after transfection with the supernatant containing MSCV-*MycN* or MSCV-vector, the C3H/10T1/2 cells were labelled with CFSE (10 μM, Abcam, ab113853) staining and seeded on 100 mm culture dishes. Two days after CFSE labeling, the cells were harvested with 0.25% Typsin-EDTA and resuspended in FACS buffer (5% FBS in PBS). Then, the cells were incubated for 15 min with DAPI (BD, 564907) in FACS buffer. The cells were analyzed by FACS Canto II (BD Biosciences) and the acquired data were further evaluated by using FlowJo software (Treestar) to calculate Mean FI. Dead cells were excluded using DAPI (Sigma-Aldrich) staining and cell doublets were systematically excluded.

### Statistical analysis

Animals from different cages, but within the same experimental group, were selected to assure randomization. Experiments involving in vitro and in vitro study was assured randomization through double-blind experiments. Data are presented as mean ± standard deviation (SD). Statistical differences between the means were compared by the two-tailed, unpaired t test for two groups, or determined using one-way ANOVA followed by Dunnett’s multiple comparison test for multiple groups. Statistical analysis was performed with Prism 9.0 (GraphPad). Statistical significance was set to P value < 0.05.

## Author contributions

S.C, and S.H.S. designed and performed the experiments, analyzed and interpreted the data; K.P.K prepared and analyzed the human samples. S.C, K.P.K. and G.Y.K. wrote and edited the paper; H.J.L contributed to *in vivo* and *in vitro* experiments; H.B. performed RiboTag mRNA-sequencing and analysis. H.J.L. and G.Y.K. directed and supervised the project.

## Acknowledgements

We thank Tae Chang Yang, Sujin Seo, and Joon-Ho Jeong for technical assistances. This study was supported by the Institute for Basic Science funded by the Ministry of Science, ICT and Future planning, Korea (IBS-R025-D1-2015 to G.Y.K.).

