## Supplemental Figures 1-3 for "Hyperactivation of YAP/TAZ drives alterations in mesangial cells through stabilization of N-MYC in diabetic nephropathy"

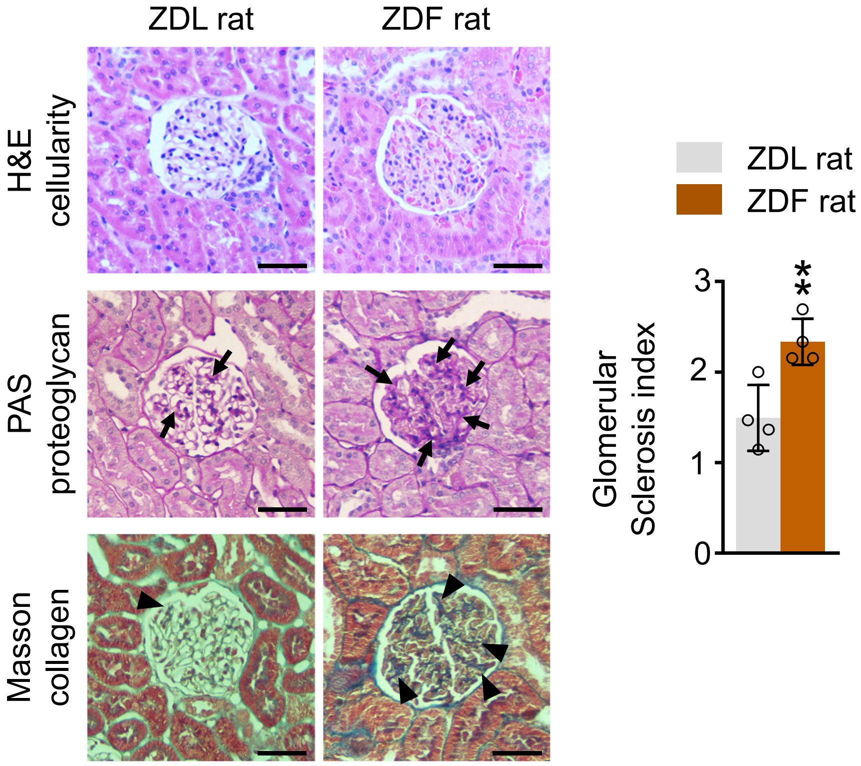


**Supplemental Figure 1. Glomerular sclerosis is occurred in the kidney of ZDF rats**

Representative images of H&E stained cellularity, PAS+ proteoglycan and Masson's Trichrome stained collagen in the glomeruli and comparisons of glomerular sclerosis index between ZDL and ZDF rats. Black arrows indicate PAS+ proteoglycan and black arrowheads indicate Masson's trichrome stained collagen. Scale bars, 50 μm. Each dot indicates a value from one rat and n= 4 rats/group from two independent experiments. Vertical bars indicate mean ± SD. **P<0.01 *versus* ZDL rats by unpaired t-test.

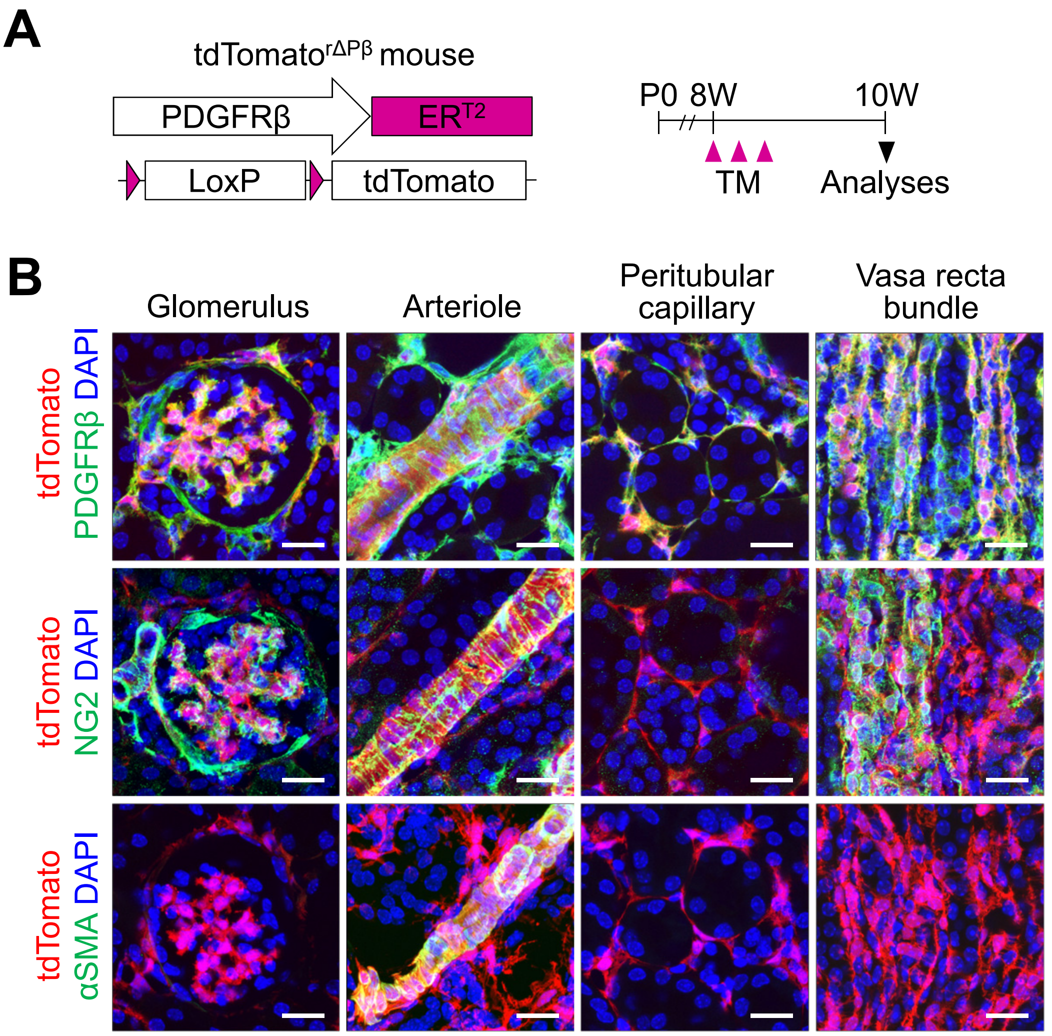


**Supplemental Figure 2. Distribution of PDGFRβ$^+$ cells in kidney**
(**A**) Diagram depicting generation of tdTomato$^{r\Delta P\beta}$ mouse and PDGFRβ$^+$ cell-specific expression of tdTomato from 8-week-old by TM administration and their analyses at 10-week-old
(**B**) Representative images of PDGFRβ$^+$ cells in the indicated region of kidney in tdTomato$^{r\Delta P\beta}$ reporter mice. Scale bars, 20 μm.

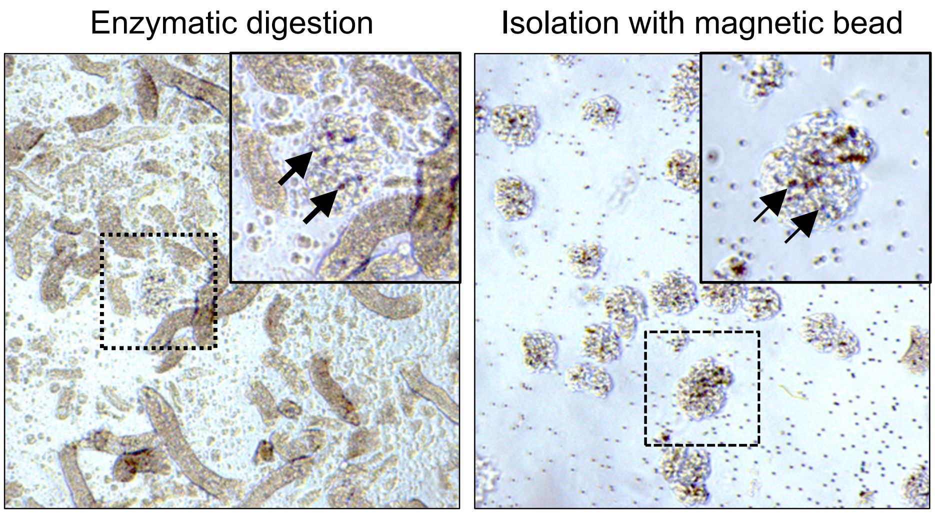


**Supplemental Figure 3. Isolation of glomeruli from the mice**
Representative images of whole kidney lysates after enzymatic digestion (left) and of glomeruli after isolation using the magnetic beads (right). Inserts are magnified views of dashed-line boxes. Black arrows indicate glomeruli with the magnetic beads (darkish brown).
